# Immune dynamics in a time of covid

**DOI:** 10.1101/2021.06.01.446677

**Authors:** Troy Shinbrot

## Abstract

Motivated by curiosities of disease progression seen in the coronavirus pandemic, we analyze a minimalist predator-prey model for the immune system (predator) competing against a pathogen (prey). We find that the mathematical model alone accounts for numerous paradoxical behaviors observed in this and other infections. These include why an exponentially growing pathogen requires an exposure threshold to take hold, how chronic and recurrent infections can arise, and what can allow very sick patients to recover, while healthier patients succumb. We also examine the distinct dynamical roles that specific, “innate,” and nonspecific, “adaptive,” immunity play, and we describe mathematical effects of infection history on prognosis. Finally, we briefly discuss predictions for some of the effects of timing and strengths of antibiotics or immunomodulatory agents.

## Introduction

Coronavirus infections provide examples of several curious behaviors that highlight the remarkable dynamics of the immune system. Some of these can be explained through a basic understanding of how the immune system confronts an invading pathogen; others cannot. For example, it is not obvious why pathogens that grow exponentially only take hold if exposure exceeds a critical value^1^. Nor is it clear why seemingly similar patients with comparable viral loads produce widely diverging outcomes, e.g. one asymptomatic and another deathly ill^2,3^. Nor is it known why some patients steadily improve after initial infection, while others worsen, improve, and then relapse^4^, sometimes repeatedly^5,6^. We examine a simple model that accounts for these and other enigmatic dynamical phenomena.

We base our model on a straightforward predator-prey framework, in which immune agents (predators: *I*_*a*_) attack pathogens (prey: *P*). Predator-prey models for immune response have been described before^7,8,9,10^; here we use these models to examine in immune dynamics in detail. We consider a version in which *P* and *I*_*a*_ change in time, *’*, according to the schematic prescription:

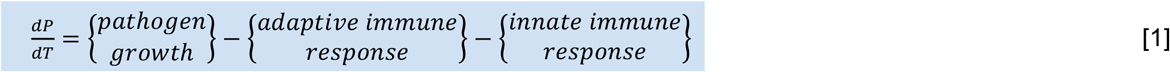

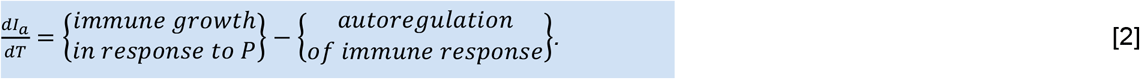

This in short says that pathogens reproduce, but are depleted by both adaptive and innate immune responses, and the immune system grows in response, but is self regulated by negative feedback. Most of these terms are straightforward to assign, as we describe next.

Before we do so, we emphasize that what we present is only a simplified mathematical model: it defines the dynamics of pathogen growth competing against idealized innate and adaptive immunity, but entirely neglects real and important biological effects known to produce complicated disease behaviors. For example, interactions between immune function and inflammation^11^, as well as autoimmune disorders^12^, are topics in their own right. Many pathogens, moreover, have evolved devious mechanisms to battle the immune system. Tuberculosis, for example, defeats a key intracellular process so that it can hijack and reproduce within the very cells that destroy most invaders^13^. Helicobacter pylori constructs a robust biofilm to isolate itself from harsh environmental and immunological attack^14,15^. The malaria parasite creates a vacuole on the membrane of liver cells, where it quietly reproduces insulated from the immune system so that it can periodically disgorge broods of new infective cells^16,17^. And HIV relentlessly mutates over the course of years until it has amassed an army of different strains that overwhelm the immune system^18^.

All of these effects produce complex disease dynamics, none of which are included in the present study. The value of the model that we present is not that it comprehensively describes the biology of pathogens and immune responses, but rather that it describes the mathematically necessary dynamics that must be present with or without other more elaborate biological effects. As we will see, these basic ingredients are sufficient to account for many curiosities seen in coronavirus and other infections.

## Model definition

### Pathogen kinetics

Pathogens typically reproduce at fixed intervals, so in Eq. [1] we set:

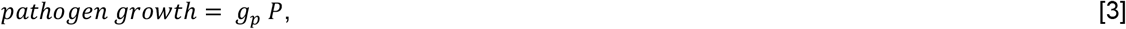

where *g* is a fixed growth rate. This produces growth defined by 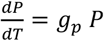, with exponential solution 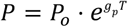, where *P*_o_ is the initial pathogen load. As for the remaining terms in Eq. [1], the immune system involves numerous interacting chemical and biological agents that are categorized in terms of whether they are adaptive (specific to a particular pathogen), or innate (nonspecific). For simplicity, we describe the adaptive response as a first order interaction, meaning a single immune cell destroys a single pathogen at a fixed rate, *k*_*a*_, and we describe the innate response to be zeroth order, meaning it depletes any invading pathogen at a fixed rate, *k*_*i*_:

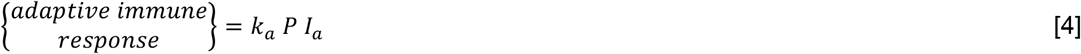

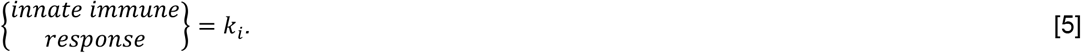

This completes the definition of pathogen growth:

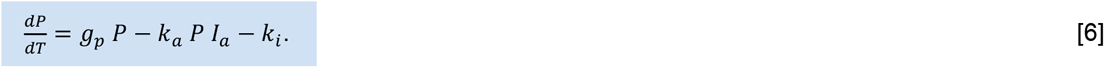

As we have emphasized already, this is by no means a comprehensive description of realistic immune response, but as a simplified model we will see shortly that it captures the essential effects of these two immune mechanisms. For a more thorough description of the intricate workings of the immune system, an enjoyable overview can be found in Ref. [19].

### Immune kinetics

As for pathogen growth described by Eq. [2], the adaptive immune system both trains cells to recognize new pathogens and recruits cells to attack known pathogens (the innate system does so as well, but to a more limited extent that we neglect here). Two central features characterize the adaptive response to pathogens: first, both training and recruitment take time, typically several days, and second both responses are “cooperative” in the biological sense, meaning that a mild infection provokes a mild response, but as the infective load grows, the immune response ramps up dramatically. Cooperative responses are often described by a “Hill” equation^20^,

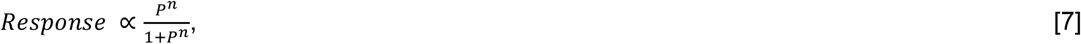

where the exponent *n* defines how cooperative the system is – i.e. how rapidly the response grows with increases in *P*. A similar response has been used previously to model white blood cell production, which of course is central to adaptive immunity^20^. So we define the growth in immune response to obey:

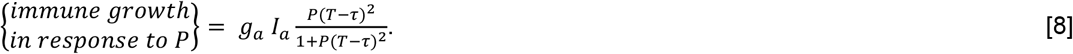

Here *g*_*a*_ is a constant defining the rate of immune growth, and the *I*_*a*_ term reflects the fact that immune cells themselves are required to produce that growth. We use *n* = *2*, which is most common in cooperative systems, but for the most part (an exception is mentioned at the end of our analysis) changes to *n* affect our results little. The Hill term explicitly depends on the pathogen load delayed by the time *τ* needed to train or recruit a response. We would expect a recently encountered pathogen to produce a rapid response (small *τ*); a novel pathogen would require a longer training time (large *τ*).

Finally, once a pathogen has been removed, the immune system returns to a quiescent state, so we include an autoregulatory term:

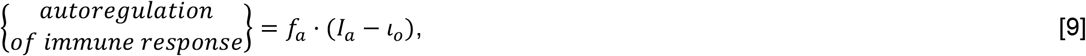

where the constant *f*_*a*_ defines the speed of return to a number, *l*_o_, of adaptive immune cells. This completes the description of the immune response:

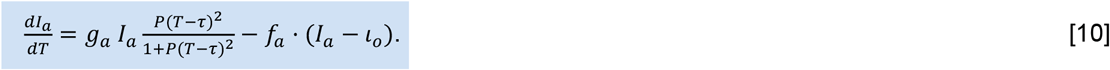

## Analysis of behaviors

Armed with Equations [6] and [10], we can now analyze how the model immune system can be expected to behave. We begin with the simplest cases and add complications sequentially.

### Innate immunity alone

The simplest two cases are (1) a system lacking any immune function, followed by (2) innate immune response alone. We examine these cases by setting *k*_*a*_ = *g*_*a*_ = 0, which eliminates adaptive immune response, and for illustration we choose representative parameters *g*_*p*_ = *2, f*_*a*_ = 1, *l*_o_ = 0.5. This transforms Eq’s [6] & [10] to:

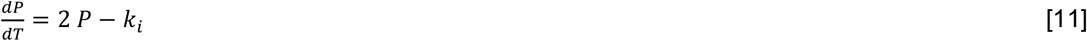

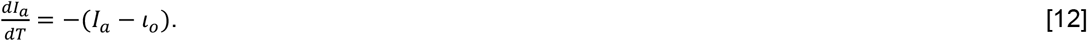

In this case, the pathogen and immune variables are independent of one another, and the equations can be solved separately, yielding two exponentials:

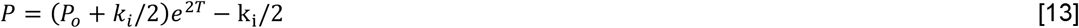

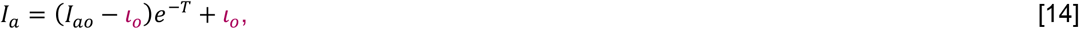

where *P*_o_ and *I*_*ao*_ are respectively initial values (numbers or concentrations) of pathogen and adaptive immune agents.

It is useful to plot trajectories for these two first cases as shown in Fig. 1 along with their nullclines^21^: curves where 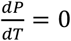 and 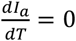. Nullclines are easily calculated from Eq’s [13] & [14] to be:

**Figure 1.**
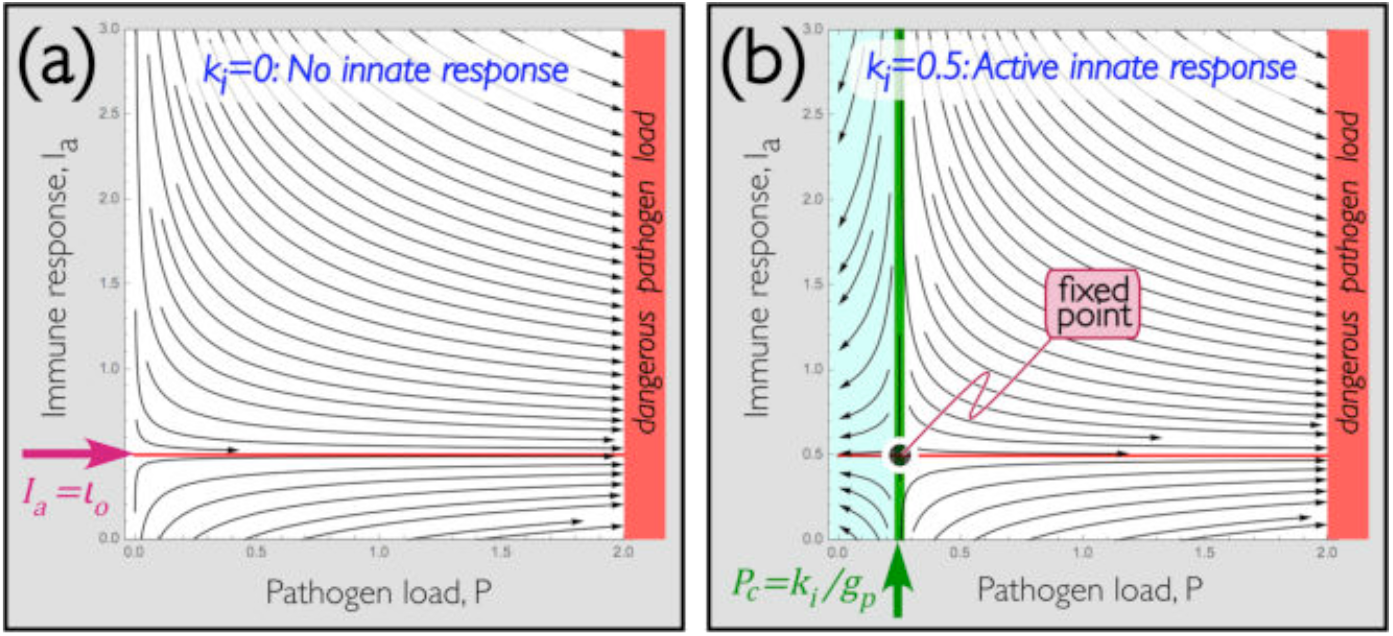
Effect of innate response alone: Trajectories for dynamics with (a) no immune response, and (b) only innate immunity. In both plots, k_a_ = g_a_ = 0, so there is no adaptive immunity. **(a)** <u>Trajectories without any immune response</u>, and with 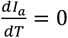 nullcline in red. As expected, trajectories have no vertical component (meaning no adaptive immune response) on this nullcline. Notice all trajectories lead to unbounded pathogen growth. **(b)** <u>Trajectories with only innate response</u>, including both earlier nullcline and 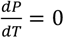 nullcline in green. There is no horizontal component on the green nullcline. Notice that in panel (a), with no immune response, pathogens grow for any initial P or I_a_, and in panel (b) with only innate immunity, a critical pathogen load P_c_ emerges (the green nullcline), below which the pathogen can be defeated. Initial pathogen loads within aqua region are cured; those within white region lead to unbounded pathogen growth. Intersections between the nullclines represent fixed points where neither P nor I_a_ change.

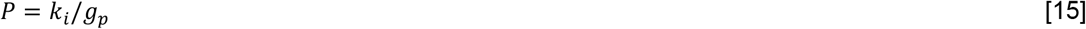

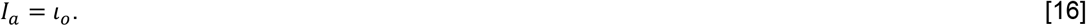

Since 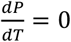 along the nullcline defined by Eq. [15], trajectories are entirely vertical along that curve (green in the figure), and similarly trajectories are entirely horizontal along nullcline [16] (red). All simulations in this work are performed in Mathematica, using its StreamPlot and NDSolve functions.

From Fig. 1(a), we can conclude, logically enough, that lacking all immune function any level of pathogen will grow – and from Eq. [13], the growth will be exponential in time. Without loss of generality, we arbitrarily choose *P* › *2* as a dangerous pathogen load, indicated by red shading in the figure. Adaptive immunity has no effect on the pathogen since we have set *k*_*a*_ = *g*_*i*_ = 0 in this example. Nevertheless trajectories approach a stable immune response at *I*_*a*_ = *l*_o_, meaning that once interactions with pathogens have been eliminated, the last term in Eq. [10] does successfully autoregulate the concentration of immune cells so that it always reaches *l*_w_.

Fig. 1(b), on the other hand, shows two things. First, the intersection between the green and red nullclines indicated is a point where both 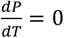 and 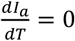, so neither *P* nor *I*_*a*_ changes there. We call this a fixed point. Second, the green, 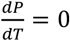, nullcline separates pathogen loads that exponentially decrease (to the left) from those that exponentially increase (to the right). This means that **the innate term produces a critical pathogen load, *P***_**c**_ = ***k***_***i***_/***g***_***p***_, **below which the pathogen will die out, and above which the pathogen will grow**.

We will see this effect in more complicated examples in following sections. Already these simplest two examples provide an explanation for a first curiosity of immune function: even though infections typically grow exponentially, steady, nonspecific depletion of pathogens is sufficient to overcome minor infections. Moreover, the size of the infection that can be tolerated is determined by the ratio [15] of the rate of innate depletion, *k*_*i*_, to the rate of pathogen growth, *g*_*p*_. For this reason, we can tolerate a relatively large pathogen load of slowly growing pathogens, but rapidly growing pathogens reduce the critical pathogen load that can be destroyed. **This is why we can tolerate limited exposure to Covid and other infections, whether we have previously been exposed to them or not**. Eq. [15] also specifies that the limit to this exposure diminishes in a predictable way as the infection becomes more virulent.

We turn next to examining how the adaptive part of Eq’s [6] & [10] behaves, considering first the adaptive system by itself, and then considering the adaptive and innate systems combined.

### Adaptive immunity alone

The innate system is defined by a single term, *k*_*i*_, in Eq. [6]. Setting that to zero and using representative values *g*_*p*_ = *2, g*_*a*_ = 1 transforms Eq’s [6] & [10] to:

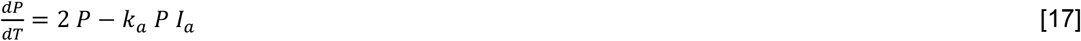

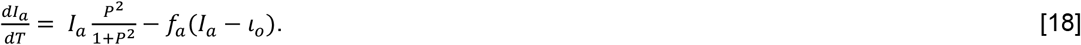

For the time being, we set the adaptive setpoint *l*_w_ = 0.5; we also neglect the effect of the time delay, *τ*, needed to train and recruit adaptive cells: we will examine effects of these terms later.

As before, it is useful to plot the nullclines, which provide a framework to understand the competing behaviors of pathogen and immune systems. Again, we do this by simply setting Eq’s [17] and [18] equal to zero and solving:

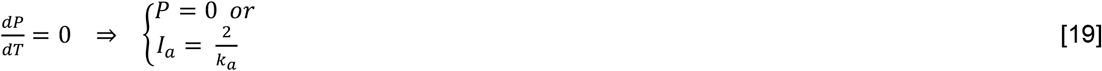

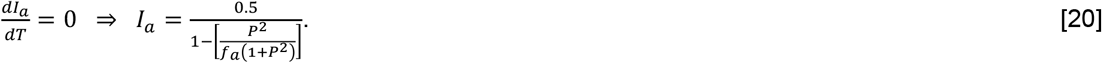

In Fig. 2, we plot these nullclines along with typical trajectories for two different values of *f*_*a*_, the autoregulatory rate at which adaptive immune cells return to the nominal setpoint, *I*_*a*_ = *l*_o_. Notice that return to the setpoint can only fully occur once *P* = 0, otherwise the first term in Eq. [18] will cause growth in *I*_*a*_. Notice also that for smaller initial adaptive immune response (*I*_*a*_‹ *l*_o_), any infection will become worse before it becomes better. We will see that this is not always the case: some infections (e.g. those above the green nullcline, which have ample immune protection) only improve, while others improve and then worsen: we will introduce these later.

**Figure 2.**
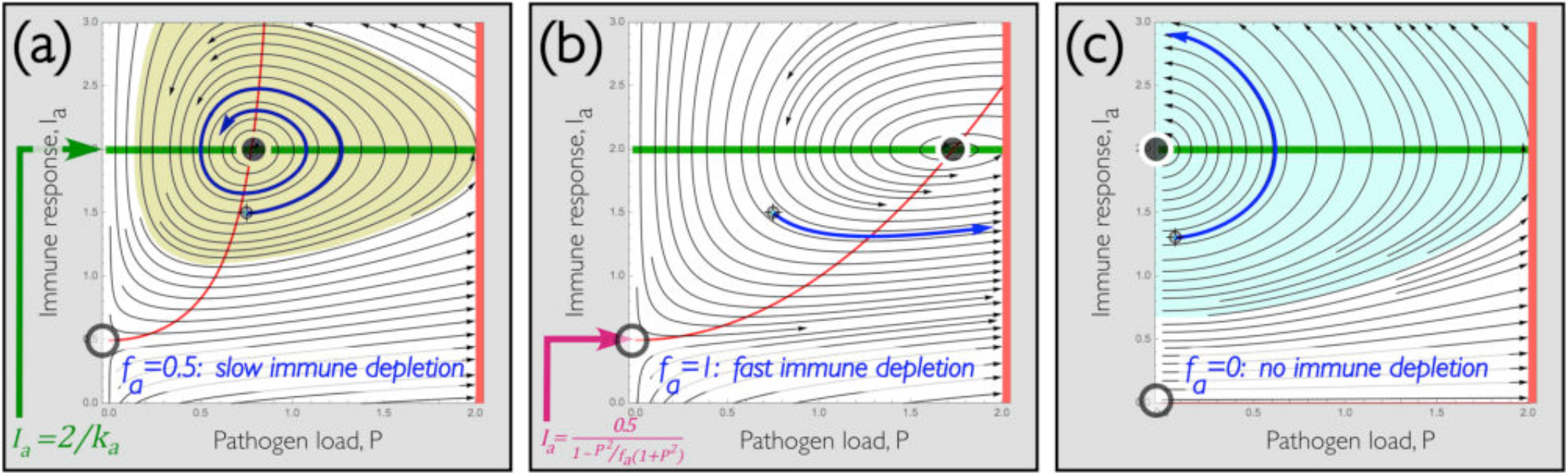
Effect of adaptive immune response alone: Trajectories showing effect of competition between adaptive immune response and pathogen load. **(a)** <u>Slow autoregulatory depletion of immune agents</u> produces inward spiral to fixed point (solid circle). Since trajectories never reach zero pathogen load, this attractive point can be interpreted as a stable, chronic, state of infection. The oscillation of trajectories around this chronic state can be interpreted as recurrent worsening and improvement of the infection. Points leading to this state are shaded sickly green. **(b)** <u>Increasing the rate of autoregulatory depletion of immune agents</u> can reduce their number faster than the growth of pathogen numbers, leading to a net worsening, both of the chronic state and of individual trajectories. Hence the sample blue trajectory, starting from the same state in both panels, reaches much higher pathogen levels – meaning that rapid immune suppression can produce unchecked infective growth. **(c)** <u>Without any autoregulation of immune levels</u>, for completeness we show that trajectories all eventually reach P = 0, but the immune level, I_a_ remains at its maximum value at that point: far above the level needed to defeat future infections. States in the aqua region are ultimately cured; those in the white region reach dangerous pathogen loads.

From examination of Eq’s [19] & [20], one fixed point is *P* = 0, *I*_*a*_ = *l*_o_, indicated by an open black circle in Fig. 2. A second fixed point is where the two nullclines intersect, as we saw in Fig. 1(b)). We identify this fixed point by a solid circle, and behavior near this point plays an important role in determining outcomes of infections. In this case, notice in Fig. 2(a) that the sample trajectory highlighted in blue spirals inward toward the fixed point. Because trajectories near the fixed point travel continually inward, the fixed point is stable: any disturbance will return the state to the fixed point. This fixed point occurs at a nonzero pathogen load, so the model apparently describes a chronic condition in which initial extremes of pathogen concentration diminish, but the pathogen is never fully destroyed. **Thus the presence of chronic infections can also be understood using this model**. At its heart, the chronic state arises because growth of the pathogen competes against depletion by adaptive immunity, but since that immunity diminishes with pathogen load, below a critical load the immune system becomes too weak to defeat the pathogen. We’ll see shortly how the model nevertheless can describe eradication of a chronic infection.

Additionally, because the blue trajectory in Fig. 2(a) repeatedly spirals around the fixed point, it represents recurrence of increased and decreased pathogen load. Many diseases have recurrences with well-identified causes – for example tertian malaria is associated with a 48-hour rhythm produced by the reproductive cycle of the underlying parasitic infection^17^. Distinct from such rhythms caused by the pathogen, there are rhythms caused by the immune system: for example fevers commonly rise and fall daily in sync with circadian physiology. We emphasize that the recurrent spiral shown in Fig. 2(a) is independent of rhythms imposed by pathogen, fever or normal physiological rhythms: it is intrinsic to the competition defined by Eq’s [6] & [10]. **So cyclic and recurrent infections are also expected outcomes from our model**.

### Effect of changes to autoregulatory rate

In Fig’s 2(b) & (c) we also display the effect of changes to immune autoregulation. Fig. 2(b) shows that an increase in the autoregulatory depletion rate, *f*_*a*_, of adaptive immune agents leaves qualitative behaviors of the pathogen-immune competition intact, but increases the chronic pathogen level – that is, the fixed point indicated by the solid circle moves to the right from Fig. 2(a) to (b). Trajectories spiral around that fixed point as before, but along the way reach much higher pathogen concentrations. So if we again say that pathogen loads beyond the right side of the plots shown are life-threatening, then the blue trajectory at the slower immune depletion rate of Fig. 2(a) would represent undesirable, but tolerable, disease levels, while the faster depletion in Fig. 2(b) would, starting from the identical initial state, produce a dangerous infection. **This means that rapidly suppressing the immune system, e**.**g. by abrupt pharmaceutical intervention, can lead to dramatic worsening in disease progression**. The root cause of this is that our model dynamics have an intrinsic time scale, and holding the pathogen in check relies on the enduring presence of immune agents. Reducing their effect too rapidly can allow the pathogen to grow too fast to be brought under control.

Contrariwise, Fig. 2(c) shows that lowering the rate, *f*_*a*_, of immune autoregulation can be expected to reduce illness and lower the chronic pathogen load. In that figure, we plot trajectories for the extreme case without any autoregulation. In that case, all trajectories end where they hit the vertical axis, i.e. at *P* = 0, *I*_*a*_ › *l*_o_. Immune agents remain at this unregulated level forever, and so the organism must tolerate an elevated immune level for the rest of its existence (a metabolic expense that evolution has presumably optimized).

### Effect of changes to autoregulatory setpoint

To recap, immune autoregulation in our model prescribes that adaptive immune levels will tend to return at rate *f* to the setpoint *l* . In Fig. 2 we considered how the rate *f* affects the system; in Fig. 3(a) & (b), we overview effects of the setpoint. In Fig. 3(a) we plot trajectories with the immune setpoint reduced, to *l*_w_ = 0, – i.e. where immune cells can respond to an infection, but are afterward autoregulated to a state with zero residual cells. This case is instructive because it is conservative, which implies that trajectories always travel in closed orbits.

**Figure 3.**
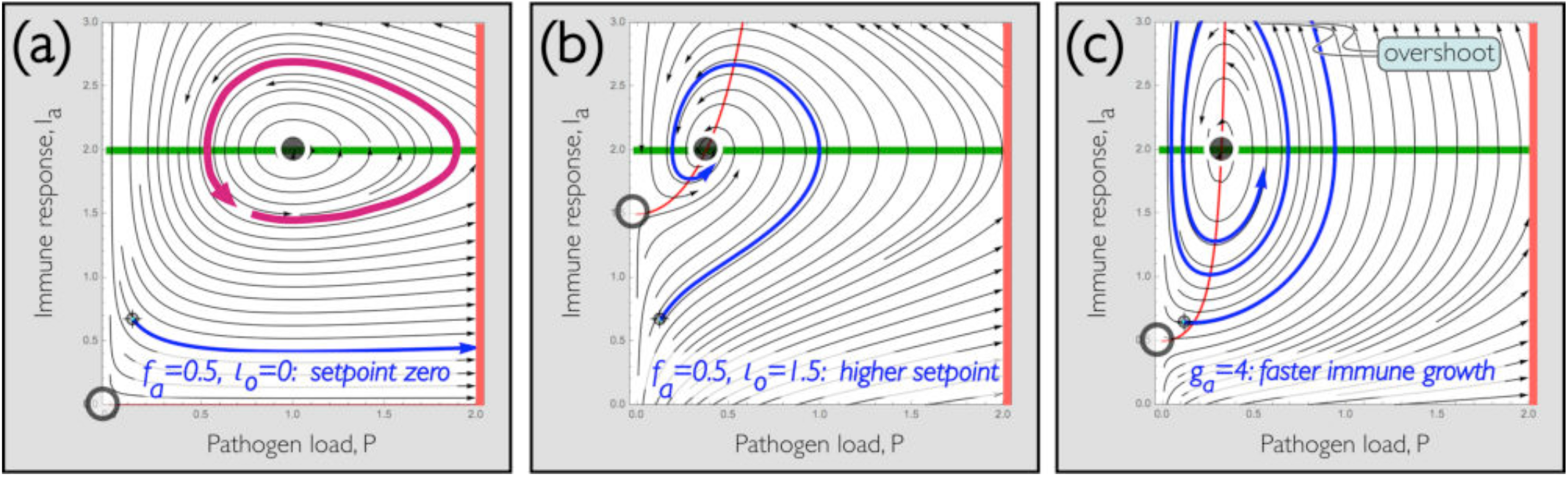
Effects of immune setpoint and growth rate: **(a)** <u>Reducing the homeostatic immune setpoint</u> to l_w_ = 0 (no residual immune agents following an infection) produces a conservative system, in which trajectories periodically encircle the fixed point, as in the magenta curve. Other initial states also encircle this point, but in concentric rings, the outermost of which that can reach dangerous pathogen levels, as shown by the blue trajectory. **(b)** <u>Increasing the number of residual immune cells</u> drives the same initial state to rapidly approach the fixed point, avoiding high pathogen levels. **(c)** <u>Increasing the rate of immune growth</u> does the same, but can produce large excursions in immune levels (overshoot indicated) and longer times to reach a steady state (panel (c) uses g_p_ = 4, f_a_ = 0.5, l_0_ = 0.5).

As an example, the magenta trajectory in Fig. 3(a) will periodically cycle around the fixed point *P* = 1, *I*_*a*_ = *2* forever. In fact almost all trajectories, for any initial state, encircle that fixed point. The blue trajectory, starting at low pathogen load and finite immune response, also does so, but it produces very large pathogen loads on its way in an extensive voyage around the fixed point. Thus just as in Fig. 2(b), where immune cells were rapidly depleted, reducing the setpoint suppresses the immune response and allows pathogen levels to grow without significant opposition.

Fig. 3(b), however, shows that unlike an increase in the rate of immune depletion, an increase in the setpoint strengthens the stabilization of the fixed point, causing the immune response to grow faster than the pathogen. As a consequence, starting from the same initial state as in Fig. 3(a), rather than producing a dangerous infective growth, the blue trajectory now shows a rapid approach to the chronic fixed point.

The implication of Fig’s 3(a) & (b) is that decreasing the immune setpoint leaves the organism vulnerable to dangerous growth of small infections. Comparing this case with Fig. 2(a) (slow immune depletion), we see that increasing the immune setpoint provides substantially better protection against infection than changing the autoregulatory rate. Apparently, and logically, **the presence of significant residual levels of immune agents is highly beneficial to the function of the adaptive immune system**. This is the mathematical interpretation of the value of prior exposure or of vaccination. Increasing the rate, *f*_*a*_, of autoregulatory control over homeostatic immune levels can reduce the metabolic expense of maintaining these agents, but without this expense, the agents will senesce or otherwise be cleared, allowing an incipient infection to grow to dangerous levels before the immune system has time to rally a response.

We have looked so far in this section at autoregulatory control over immune response. This is encapsulated in the last term in Eq. [10], which depends on *f*_*a*_ and *l*_*o*_, but plainly the adaptive immune growth, specified by *g*_*a*_, competes against this autoregulatory depletion. To explore the effect of this competition, in Fig. 3(c) we increase that rate from its nominal value *g*_*a*_ = 1 up to a new value, *g*_*a*_ = 4: twice the rate of pathogen growth, *g*_*p*_ = *2*. The other parameters are returned to their nominal values originally shown in Fig. 2(a): *g*_*p*_ = *2, f*_*a*_ = 0.5, *l*_0_ = 0.5.

As Fig. 3(c) shows, by increasing *g*_*a*_ we achieve much the same end as increasing the immune setpoint, *l*_o_. This makes sense: one can achieve a similar goal either by starting at a higher immune level or by driving the system to that level faster. Fig. 3(c) also shows, however, that faster immune growth causes substantial overshoot, producing large excursions in immune levels, and consequently the system takes longer to reach steady state.

### Combined innate and adaptive immunity

Summarizing findings up to this point, we have seen that the ***innate*** immune system by itself can overcome minor infections. Mathematically speaking, as it does so an ***unstable*** demarcation (the green nullcline in Fig. 1(b)) emerges between tolerable and dangerous infections. The ***adaptive*** system alone can constrain larger infections, but it tends to ***stabilize*** those infections, leading to a chronic state (that may or may not oscillate depending on autoregulatory details).

The interplay between unstable and stable states is central to understanding the combination of innate and adaptive immunity, and in Fig. 4 we show several different behaviors produced by this interplay as the innate strength, *k*_*i*_, is increased.

**Figure 4.**
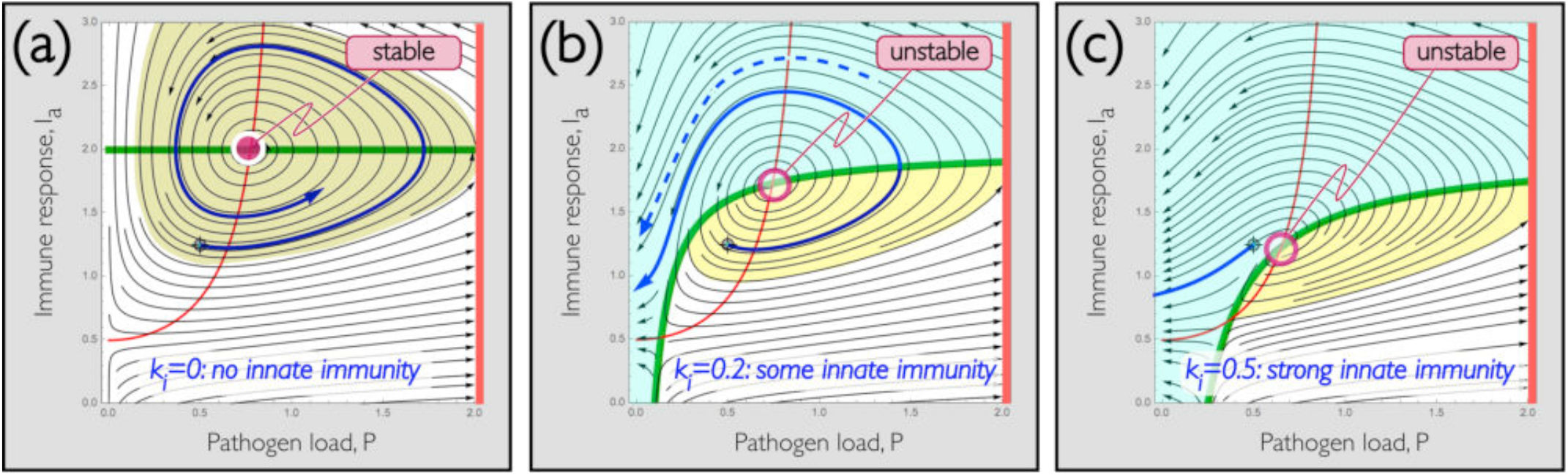
Combined innate and adaptive immunity: **(a)** <u>Copy of Fig. 2(a) with fixed point identified</u> as stable. Sickly green points are attracted to fixed point. **(b)** Under identical conditions but <u>increasing the innate immunity coefficient to k_i_ = 0.2</u> makes the fixed point unstable. As a consequence, the same initial condition that was attracted to the fixed point in panel (a) is now repelled, leading to zero pathogen load as shown in solid blue trajectory. As in prior plots, infections that start within the aqua region lead to cure; infections within the yellow region do so as well, but the disease gets worse before it gets better. Very low immunity, in white, gets steadily worse. **(c)** Modeling a <u>boost to the immune response by further increasing k_i_ to 0.5</u> causes the green nullcline to move to higher pathogen loads. This nullcline divides regions that steadily improve from those that worsen before they improve. So increasing innate immune function – i.e. increasing k_i_ – causes the same initial state as shown in solid blue in panels (a) and (b) to be rapidly cured.

Starting with Fig. 4(a), we show trajectories produced with *k*_*i*_ = 0, i.e. without innate immune function. As in the earlier Fig. 2(a), we use illustrative values *g*_*p*_ = *2, k*_*a*_ = *g*_*a*_ = 1, *τ* = 0, *f*_*a*_ = *l*_0_ = 0.5 in Eq’s [6] and [10]. In Fig. 4(a), we identify the stable fixed point that attracts nearby trajectories as a solid magenta circle, and we show a representative orbit around the fixed point in blue.

Beginning at the same initial point, Fig. 4(b) shows the solid blue trajectory in the presence of both adaptive and innate immunity, produced by setting *k*_*i*_ = 0.*2*. Notice that this trajectory still encircles the fixed point, but rather than being attracted, it is repelled by that point – hence we now identify it with an open circle using the convention introduced in Fig. 2. That is, introducing innate immunity has caused the fixed point to transition from stable (attractive) to unstable (repulsive). This occurs in this example when the innate immunity strength in Eq. [6], *k*_*i*_, exceeds about 0.1018. When the fixed point becomes unstable, it pushes the solid blue trajectory away as we see in Fig. 4(b), so that it now reaches *P* = 0 – which is to say so that all pathogens are destroyed.

**This means that combining innate and adaptive immunity allows more serious infections to be controlled, though neither immune response alone can do the job.** The innate system alone can eliminate minor infections but cannot control larger ones. The adaptive system alone can stabilize larger infections, but tends to reduce its intensity as the infection diminishes – and hence can produce chronic conditions. We will see that the delay of the adaptive system described by Eq. [10] is also important to chronic conditions, but for now we settle for the observation that combining innate and adaptive responses can effectively combat infections that neither can cure alone.

Notice that in Fig. 4(b) there are two qualitative outcomes. <u>First</u>, trajectories beginning in the shaded aqua region above and left of the green nullcline lead monotonically to decreased pathogen load – that is, outcomes steadily improve. An example is shown as a dashed blue line. Trajectories in this aqua region that begin to the left of the red nullcline see a decrease in both immune and pathogen levels; those above the green but to the right of the red nullcline require an increase in immune expression to destroy the pathogen. In either case the pathogen is steadily and completely removed.

<u>Second</u>, trajectories like the solid blue line that start in the yellow region produce an increased pathogen level before the pathogen is removed, meaning **the disease gets worse before it gets better**. We have seen this before in Fig’s 2 and 3 – we will see shortly, though, that some diseases in this model can do the opposite: i.e. get better and then get worse.

Before we examine more complicated cases, note that even stronger innate immunity drives green nullcline further to the right: this is shown in Fig. 4(b), where we raise *k*_*i*_ from 0.2 to 0.5. To stretch the point, this could also be considered to result from an adjuvant that boosts a weak immune response. The green nullcline, recall, distinguishes trajectories that monotonically improve from those that first worsen: this occurred as well in Fig. 1(b) without adaptive immunity. A companion finding with combined innate and adaptive community is that the green nullcline can move past an initial state and so switch the outcome between one that steadily improves and one that worsens. For example the solid blue trajectory in all panels of Fig. 4 starts from the same location, but leads in panel (a) to a recurrent infection, in panel (b) to an infection that worsens and then resolves, and in panel (c) to an infection that rapidly and monotonically improves. **Thus a moderate increase to model innate immunity can resolve chronic infection following a period of recurrence: a larger increase can rapidly resolve the same infection**.

### Basins of distinct immune dynamics

We showed in Fig. 4 that adjacent initial states can lead to steadily worsening (white), improving (aqua), as well as chronic or recurrent (sickly green) infections. We also mentioned that some states worsen and then improve, while others improve and then worsen. In Fig. 5(a), we identify regions, or “basins” that lead to each of these outcomes for the combined innate and adaptive immunity case shown in Fig. 4(b). Fig. 5(b) shows an enlargement near the central fixed point: states within the black basin ultimately reach the redline indicating a dangerous pathogen load (i.e. patient death). In both panels, colored regions ultimately get better (*P* → 0); colorless regions ultimately get worse *P* › *2* . The boundaries between distinct basins interleave closely, so **small differences in initial state can produce large differences in outcome**.

**Figure 5.**
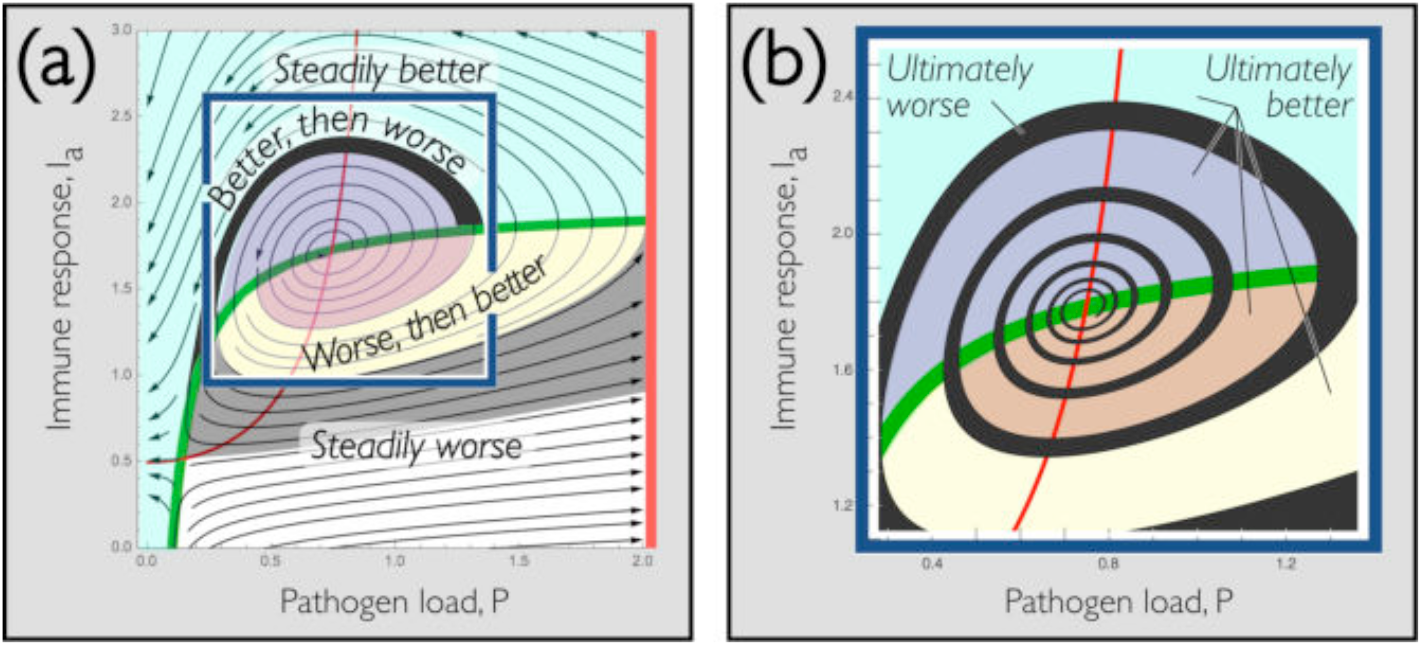
States that worsen alongside states that improve: **(a)** <u>Same conditions as Fig. 4(b), color coded based on outcome</u>. At the highest I_a_ values (aqua), essentially any pathogen load produces steady improvement. In the lower I_a_ band identified in black, the disease improves (P decreases) and then worsens (P increases). Lower I_a_ still (yellow) produces worsening then improvement, and the lowest I_a_ (gray or white) produces steady worsening. **(b)** <u>Enlargement of region near fixed point</u> shows that the outcome can ultimately worsen or improve depending on exact initial state. Initial conditions (black) that ultimately worsen spiral inward toward the fixed point. Initial conditions within colored regions ultimately improve.

### Effect of initial state

The strong dependence of outcome on initial state can be traced to the root cause, shown first in Fig. 2, that there are two fixed points in our simplified model. We highlighted the importance of stability of a first fixed point in Fig’s 4 and 5; the second fixed point is circled in Fig. 6 closer to the origin. Two nearby trajectories that approach this fixed point are shown in blue in Fig’s 6(a) and (b); both panels use identical parameters as used in Fig. 4(b). A first trajectory, in Fig. 6(a), leads to a rapid elimination of all pathogens, while a second, in Fig. 6(b), produces an exponential growth in pathogen levels. The parameters are exactly the same in both panels, but pathogen loads initially differ here by a part in 1000. This deviation can be arbitrarily small, for instance initial values of *P, I*_*a*_ = 0.3824534, *2* and 0.3824535, *2* differ by about 3 parts in 10 million (approaching the limits to computational precision), and again the first leads to a rapid cure while the second leads to dangerous pathogen growth. It is not difficult to show that the circled fixed point is always unstable^22^ in this system, and this sensitivity to initial infection load is intrinsic to systems with unstable fixed points such as this. Thus the finding that **very similar patients can experience very different outcomes is a natural expectation of the simplified model**.

**Figure 6.**
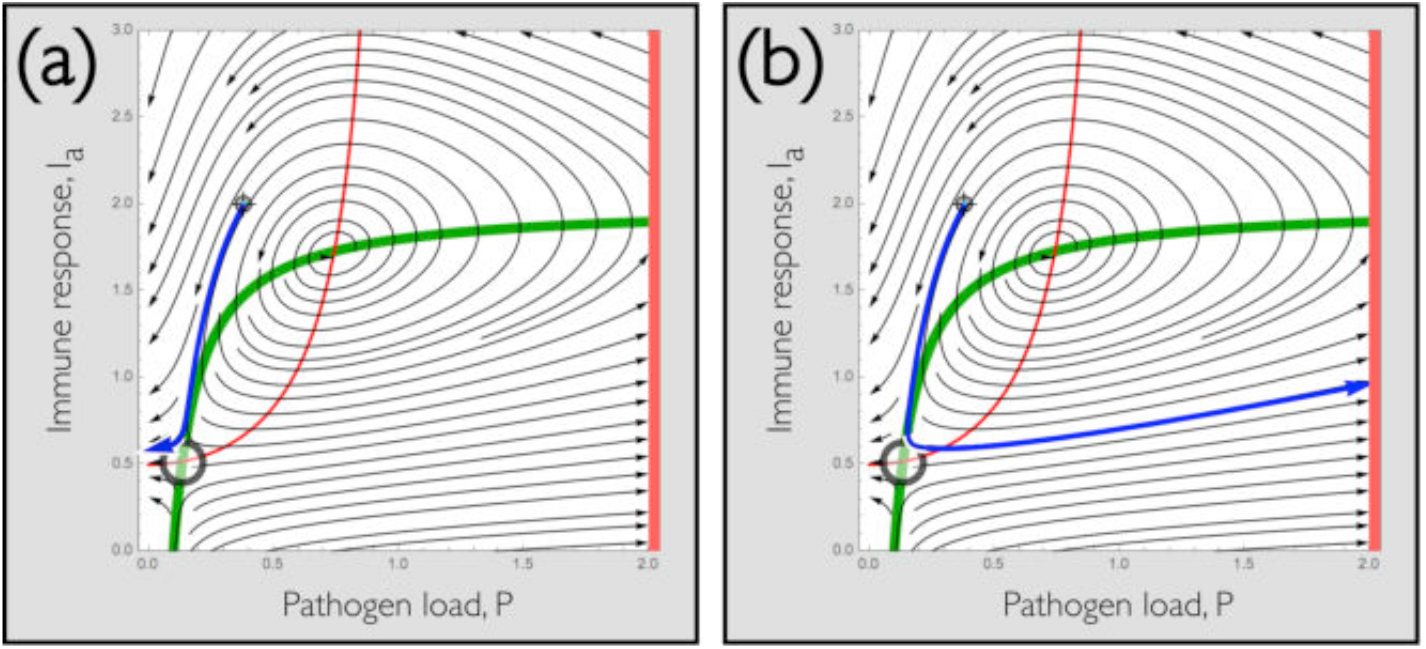
Diverging outcomes. Simulated disease progressions under identical conditions, differing only by 0.1% in initial pathogen exposure, produce very different outcomes. **(a)** <u>Blue curve uses initial state P, I_a_ = 0.3822,2</u>, which shows a rapid and complete cure. **(b)** <u>Blue curve with initial state P, I_a_ = 0.3826,2</u>, which shows exponential pathogen growth. In both panels, the same parameters are used as in Fig. 4(b): g_p_ = 2, k_a_ = g_a_ = 1, τ = 0, f_a_ = l_0_ = 0.5, k = 0.2.

### Controlling trajectories – early intervention and limits to treatment

We learned from Fig. 4(b)-(c) that changing parameters such as rates of growth or regulation can move nullclines and so alter a disease’s progression. From Fig. 5, we saw that the same can be accomplished by changing the state, i.e. the pathogen load, *P*, or immune response, *I*_*a*_. This raises the obvious question of whether outcomes can be intentionally controlled by making changes to parameters or variables. For example, effects of antibiotics can be modeled by recalling that the final, innate, term in Eq. [6] reduces infections at a fixed rate, and mathematically speaking this could just as well be caused by exogenous drug treatment as by endogenous immunity.

As a case in point, it is clinically recognized that early intervention is important in the treatment of many diseases: a fact that is commonly ascribed to causes including the reduction in inflammation, tissue damage and biofilm formation by the offending organism. In addition to these very real causes, a fundamental dynamical effect is at work, as illustrated in Fig. 7. In both panels, we plot the movement of the 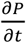 nullcline produced by an increase in *k*_*i*_ from 0.1 to 1 – representing a faster depletion of a pathogen that might be produced by an antibiotic treatment. In panel (a) we illustrate an early intervention, showing treatment when the state has reached the red star, to the left of the moved nullcline. In panel (b) we show the same treatment applied later, when the state has reached the red star at higher pathogen load. The resulting trajectories in both cases are shown in blue, and follow the underlying trajectories, both at *k*_*i*_ = 0.1 and *k*_*i*_ = 1 shown in gray. **Evidently, early intervention can produce a cure that later intervention – even using a stronger treatment – may fail to achieve**.

**Figure 7.**
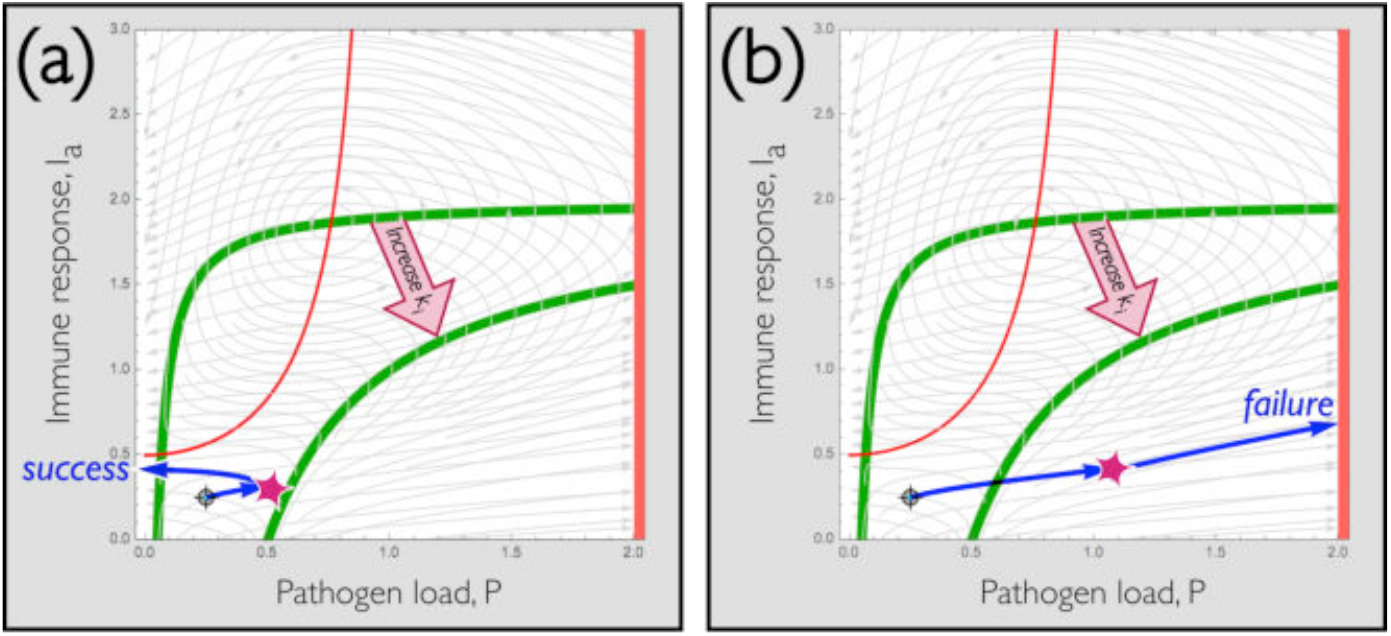
Value of early intervention: Increasing the rate of innate pathogen depletion from k_i_ = 0.1 to k_i_ = 1 lowers the green nullcline as indicated by the arrow. The moment when the increase occurs is indicated by the red star, and all trajectories under both parameter sets are shown in gray. **(a)** <u>Early intervention</u>: If k_i_is increased before the pathogen load has grown beyond a point of no return, the trajectory can change direction and the treatment will be successful as shown by the blue trajectory. **(b)** <u>Later intervention</u>: If the increase in k_i_ is delayed, the pathogen load may reach a point (again indicated by the star) beyond the new green nullcline, where all trajectories lead to dangerous growth in P, again shown as a blue trajectory. Thus some infections that can be treated successfully by early intervention can fail to be treated after the pathogen load has grown sufficiently.

### Effects of delay

An essential feature of adaptive immunity is that it takes time to train or recruit a specific immune response^20^. The delay, *τ*, is included in Eq. [10] to account for this time, however including *τ* carries a mathematical burden: in order to solve for the state at a time, *T*, one must know the history of *P*(*t*) and *I*_*a*_(*t*) at all times *t* between *T* − *τ* and *T*. Since there is an infinite number of such times, such “delay differential equations” are formally infinite dimensional, and consequently it is not possible to provide a uniformly applicable solution.

In practice, the extent to which delay differential equations depend on history varies considerably, and the study of these equations is a topic unto itself^23^. To make numerical solutions tangible, we will consider two of the infinite number of possibilities. First, we will examine the situation in which a disease initially changes slowly compared with the time delay *τ* – that is, we define the initial pathogen history to be constant, *P t* = *P*_o_ for – *τ* ‹ *t* ‹ 0. This is a simple case that can be reasonably cleanly analyzed. Second, we will examine a pathogen that grows exponentially at the fixed rate *g*_*p*_ up to the value *P*_o_, i.e. 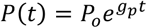 during the time – *τ* ‹ *t* ‹ 0. This case is more realistic, but since the pathogen load changes in time before the immune system has been engaged, it produces hysteretic effects that can seem paradoxical.

### Constant history

We start by considering the case shown in Fig. 4(c), with strong innate immunity (*k*_*i*_ = 0.5), and the other parameters as before: *g*_*p*_ = *2, k*_*a*_ = *g*_*a*_ = 1, *τ* = 0, *f*_*a*_ = *l*_0_ = 0.5. Without no delay, *τ* = 0, we reproduce the earlier results in Fig. 8(a) and superimpose a new representative trajectory in blue. The initial state shown produces a worsening of infection followed by a reduction in both *P* and *I*_*a*_. This is a satisfactory outcome: the pathogen is eliminated, and autoregulation reduces the immune response to close to its original level.

**Figure 8.**
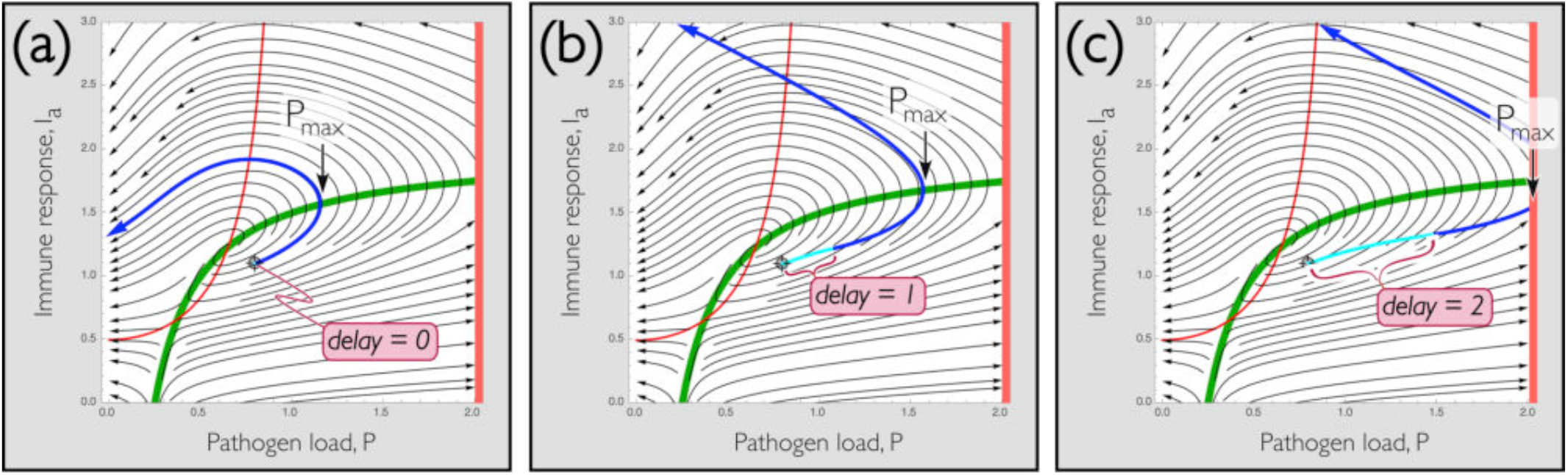
Effect of delay: Increasing the delay in Eq. [10] allows the pathogen load to increase while not increasing the immune response. **(a)** <u>No delay</u>: Without delay, all trajectories follow streamlines generated in earlier plots. **(b)** <u>Modest delay</u>: Immune response is retarded, causing both the maximum pathogen load, P_max_, and the ultimate immune response (not shown) to grow. Cyan indicates the portion of the trajectory computed using fixed P and I_a_ during the delay-time before the simulation begins. **(c)** <u>Long delay</u>: Increasing the delay results in a longer period (cyan) during which the immune response is prescribed and does not yet respond to the current pathogen load. This permits the pathogen load to continue to grow without response, leading to a larger P_max_ (here beyond the dangerous redline).

As the delay is increased, the immune response is further retarded, which allows the pathogen to grow for a time before the adaptive system responds. As a result, the maximum pathogen load, *P*_*max*_ in Fig. 8, increases with delay time. Fig’s 8(b) & (c) show trajectories starting from the same initial state as in Fig. 8(a), but with delays *τ* = 1 and *τ* = *2* respectively. At *τ* = 1 (Fig. 8(b)), *P*_*max*_ increases modestly; more significantly the adaptive immune response begins later in the game, and so the ultimate value of *I*_*a*_ at which the pathogen is destroyed is much larger than before. This is beneficial from the standpoint that the immune system ends up highly energized and ready to combat a similar infection, but also potentially results in over-activation of the immune system. To make clear what the immune-pathogen competition is doing both in the “pre-history” stage, before the simulation begins at *t* = 0 and afterward, we color the trajectory cyan during the period when the initial pathogen is held constant (– *τ* ‹ *t* ‹ 0), and blue thereafter *t* › 0 . The cyan part of the trajectory depends on the prior history; the subsequent trajectory in blue is reliable given the history shown in cyan. We will describe some aspects of how history can affect outcome shortly.

At longer delay yet, *τ* = *2* in Fig. 8(c), *P*_*max*_ grows beyond the redline, indicating a dangerous pathogen load. This is to be expected: essentially the immune system is retarded beyond the time when it is needed, so *P* can grow almost unconstrained. This fact allows us to redefine the basin of attraction for states that can be cured as a function of delay.

So for example, recall from earlier figures (e.g. Fig. 4) that we identified initial states in yellow and aqua that lead to a cure (*P* = 0). As a delay is introduced, the curable region contracts as shown in Fig. 9 for delays ranging from *τ* = 0 to *τ* = 4. In Fig. 9(a), we show the example of strong innate immunity, *k*_*a*_ = 0.5, earlier considered in Fig. 4(c). Initial states in the cross-hatched area would be curable with no delay, but as delay is introduced, the curable area becomes eroded – meaning that fewer states would be curable.

**Figure 9.**
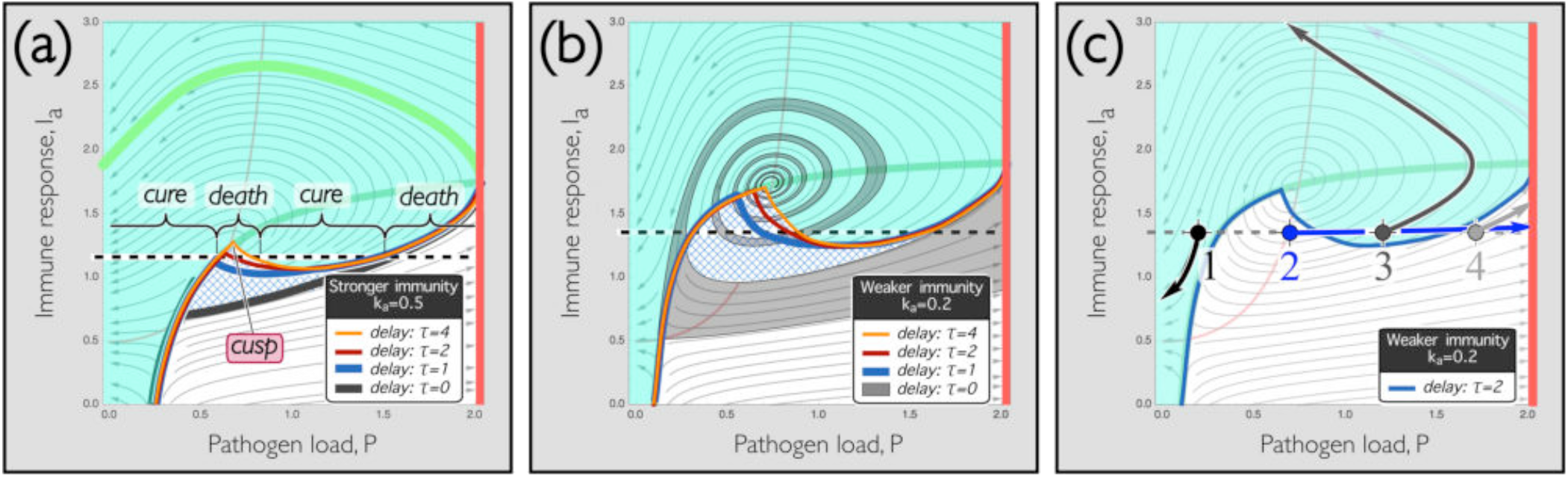
Cusp catastrophe permits pathogen to sneak up on immune system: Aqua and cross-hatched regions indicate initial states that ultimately lead to a cure. **(a)** <u>Stronger immunity</u>: Using k_a_ = 0.5 (cf. Fig. 4(c)), longer delays erode the cure basin: the cross-hatched region shows initial points that would be cured for τ = 0, but not for longer delays. Notice that a cusp grows as the delay increases, and so for nonzero delays, an immune response such as shown by the dashed line results, moving left to right, in a cure for small P, death for larger P, cure for still larger P, and death for the largest P values. **(b)** <u>Weaker immunity</u>: Using k_a_ = 0.2 (cf. Fig. 4(b)), the region of initial points that are cured shrinks. Note that with no (or small) delay, nearby initial states that lead to cure or death interweave as shown in Fig. 4, but is eliminated by either stronger immunity (panel (a)) or significant delay (this panel). **(c)** <u>Sample trajectories</u>: Four trajectories starting from initial points indicated, using k_a_ = 0.2, τ = 2, confirm that states evolve sequentially to **1**: cure, **2**: death; **3**: cure, and **4**: death. Notice as described in text that the delay term in Eq. [10] implies that trajectories with different histories can cross one another, which does not occur without delay, as in Fig. 4.

As one might expect, when adaptive immunity is weakened, the range of states that can be cured is diminished, and even more so with delayed response. This is shown in Fig. 9(b), where we plot basin boundaries as in 9(a), but with immunity reduced from a stronger value, *k*_*a*_ = 0.5 to a weaker one, *k*_*a*_ = 0.*2*. The cross-hatched area grows, indicating that a larger range of states is sensitive to delay when the immune system is weakened. More interestingly, a cusp develops as identified in Fig. 9(a). Cusps and their consequent “catastrophes” have been well studied^24^; the result here is that for a fixed immune response – for example, *I*_*a*_ = 1.35 as indicated by the dashed line in Fig. 9(b), increasing the pathogen load causes the ultimate outcome to switch from cure (at low *P*) to death (above about *P* = 0.3) to cure again (*P*∼0.8) and finally to death (above *P*∼1.6). The same occurs at stronger immunity (Fig. 9(a)), but the effect is less pronounced.

For any immunity strength, this is a highly counterintuitive finding: it means that although very small or very large pathogen loads logically enough lead to patient recovery or death respectively, between those extremes **one patient with a lower load may die, while an identical patient with a higher load may survive**. The cause of this paradoxical result is that a higher pathogen load can provoke a stronger immune response that, combined with delay, can lead to a successful outcome, while a lower load can permit the pathogen to gain in strength, “sneaking up” on the immune system so to speak, and ultimately overcoming it.

This may be made more evident in Fig. 9(c), where we plot trajectories starting from four initial pathogen loads, **1**: *P* = 0.*2*, **2**: *P* = 0.*7*, **3**: *P* = 1.*2*, and **4**: *P* = 1.*7*, all for the same immune response, *I*_*a*_ = 1.35. The first state is promptly cured, the second produces steady pathogen growth, the third worsens and then steadily improves, and the last rapidly dies. Notice that the trajectory labeled **2**, in blue, crosses trajectories **3** and **4** – that is, starting from state **3** (with *P* = 1.*2* and *I*_*a*_ = 1.35 held constant for time *τ* = *2*) leads to a cure. This occurs because Eq. [10]:

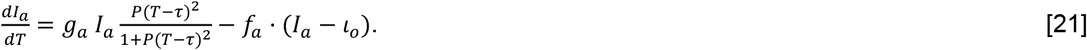

holds 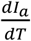 constant, at a relatively high value, for the period from time *T* = −*τ* until time *T* = 0 when the simulation begins. Meaning that *I*_*a*_ can grow at this constant value as soon as the simulation begins: as the trajectory beginning at state **3** shows. On the other hand, beginning at state **2** and passing through state **3**, the simulation is held at the relatively lower value of *P* that was established at state **2**. Indeed, state **2** starts very near to the red 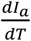 nullcline, so *I* changes by very little initially: as the trajectory beginning at state **2** shows. Thus the pathogen can grow, or sneak up on, the immune system, leaving it uninfluenced by the pathogen growth.

We remark that it may appear from Fig. 9(b) that the switching of cure and death outcomes is superficially similar either with or without delay. That is, if one travels along the dashed line in Fig 9(b) either *τ* = 0 or with *τ* = *2*, one will in both cases encounter alternating regions leading to cure and to death. Mathematically, however, despite the apparent qualitative similarity, these situations are different. Without delay, trajectories are strictly two-dimensional, because the dynamical system defined by the ordinary differential Equations [6] and [10] is itself two-dimensional, and resulting trajectories cannot cross one another. As we have remarked, however, the dynamics in the presence of delay is higher dimensional, and so trajectories in basins that lead to cure or death can cross one another (or more precisely, appear to cross in the two-dimensional projection of the underlying higher dimensional system). Crossings shown in panel (c) carry the meaning that given a state, *P, I*_*a*_, at a time *t*, different outcomes can emerge depending on the history prior to *t*.

### Exponential history

Apropos of this observation, the specific outcome shown in Fig 9 obtains in our first, simpler, choice of history in which *P* is held constant for time *τ* before beginning the simulation. A different choice of history would produce a different specific outcome. Nevertheless, it is a general result independent of choice of history that **delays allow trajectories to cross so that a smaller initial pathogen load can produce a larger ultimate infection**.

To see this, we turn to the more realistic case of a pathogen that grows exponentially in time with rate *g*_*p*_, so that 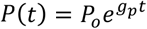 during the time – *τ* ‹ *t* ‹ 0. Not surprisingly, this more elaborate history results in more elaborate trajectories. In Fig. 10, we plot results using this exponential history with the same parameters as in Fig. 9(b). We see that the relatively modest cusps produced previously become embellished into a spiral wave shape. In detail, as delay is increased, the original spiral shown in Fig. 4 without delay (gray in Fig. 10(a)) unwinds, producing a basin boundary that bends through *π* radians as shown at around *τ* = 1 (blue), about *π*/*2* radians near *τ* = *2* (red), and becomes single valued around *τ* = 4 (orange). That is, for large delays, corresponding to each adaptive response value there is a single critical pathogen load, *P*_*c*_, above which the pathogen will grow. The emergence of a critical load is similar to the case without adaptive response, except that the value of *P*_*c*_ grows with *I*_*a*_ as shown in Fig. 10(a), and indeed the longer the delay, the more vertical (i.e. independent of *I*_*a*_) the critical load becomes. This makes sense: as the saying goes, “justice delayed is justice denied,” and in the same way delaying an adaptive immune response long enough is equivalent to removing it altogether.

**Figure 10.**
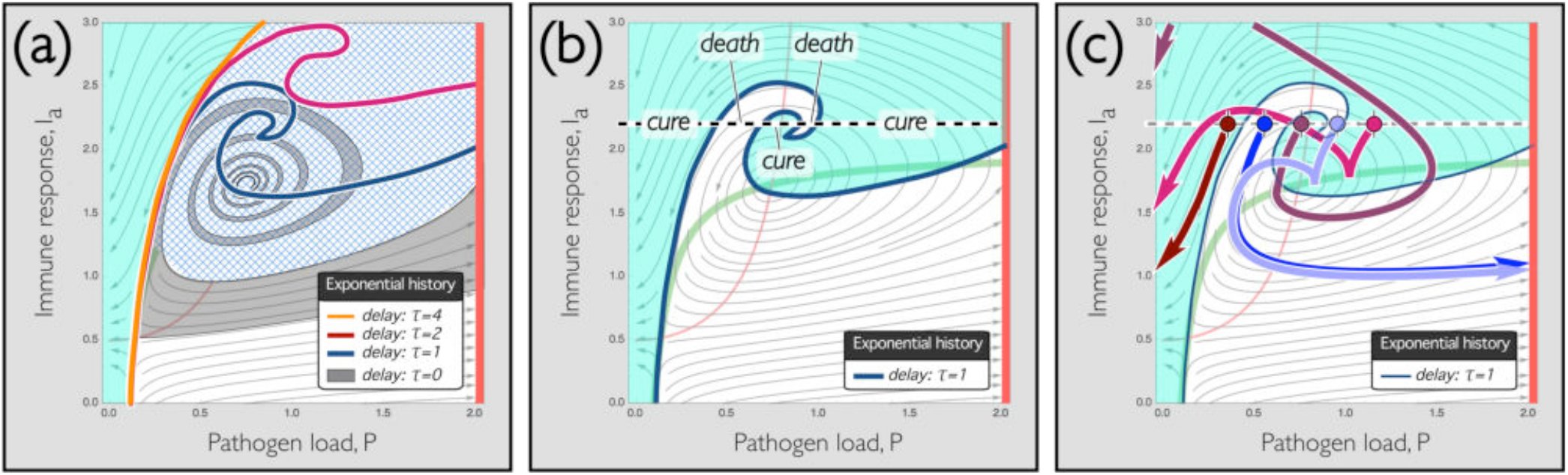
Exponential history elaborates basin boundary: If the pathogen grows exponentially (rather than remaining constant as before), the situation becomes more elaborate. **(a)** <u>Growth in dangerous basin with delay</u>: For delays τ = 0 to τ = 4, the basin of initial conditions leading to death grows substantially. As before, aqua indicates initial states that lead to cure; cross-hatching indicates initial states that would be curable without delay, but become dangerous with delays indicated. **(b)** <u>Multiple outcomes</u>: τ = 1 case – notice that for an initial immune value of I_a_ = 2.2 there are now <u>five</u> outcomes for sequentially increasing pathogen load, from left to right along the dashed line, cure, death, cure, death, cure. **(c)** <u>Sample trajectories</u>: five trajectories, all at I_a_ = 2.2 and with P = {0.35, 0.55, 0.75, 0.95, 1.15} are shown, leading alternately to cure and death as described previously.

In Fig. 10(b), we highlight the spiral wave shape for a delay *τ* = 1: evidently in this case, at a constant initial immune response *I*_*a*_ = *2*.*2*, the exponential growth of pathogen produces a sequence of five alternating outcomes. For *P* in the three ranges in aqua ([0, 0.43] or [0.68, 0.89] or [0.98, 2]) a cure results; for *P* between these ranges, in white, death results. Fig. 10(c) displays trajectories, which again appear to cross, in each of these ranges. Evidently, **both simplistic (Fig. 9) and more realistic (Fig. 10) history models produce paradoxical outcomes, so that for the same immune state a worse infection can produce a better result**.

### Limit cycles and more

We mentioned earlier that delay differential equations can be very complicated; a few examples of this are shown in Fig. 11. First, we know from Fig. 4 that increasing the innate immunity (controlled by *k*_*i*_) causes the stable fixed point to become unstable, and without a delay, we saw that this allows combined innate and adaptive immune systems to cure chronic diseases. It turns out that in the presence of delay^25^, trajectories become bounded rather than exiting the domain either below *I*_*a*_ = 0, which we designated a cure, or above *I*_*a*_ = *2*, which we associated with death. We demonstrate that trajectories become bounded in the dark blue trajectory in Fig. 11(a), which migrates inward as shown. In that panel, we use *g*_*p*_ = *2, k*_*a*_ = 1.1, *k*_l_ = 0.015, *g*_*a*_ = 1, *f*_*a*_ = *l*_0_ = 0.5 and *τ* = 0.155.

**Figure 11.**
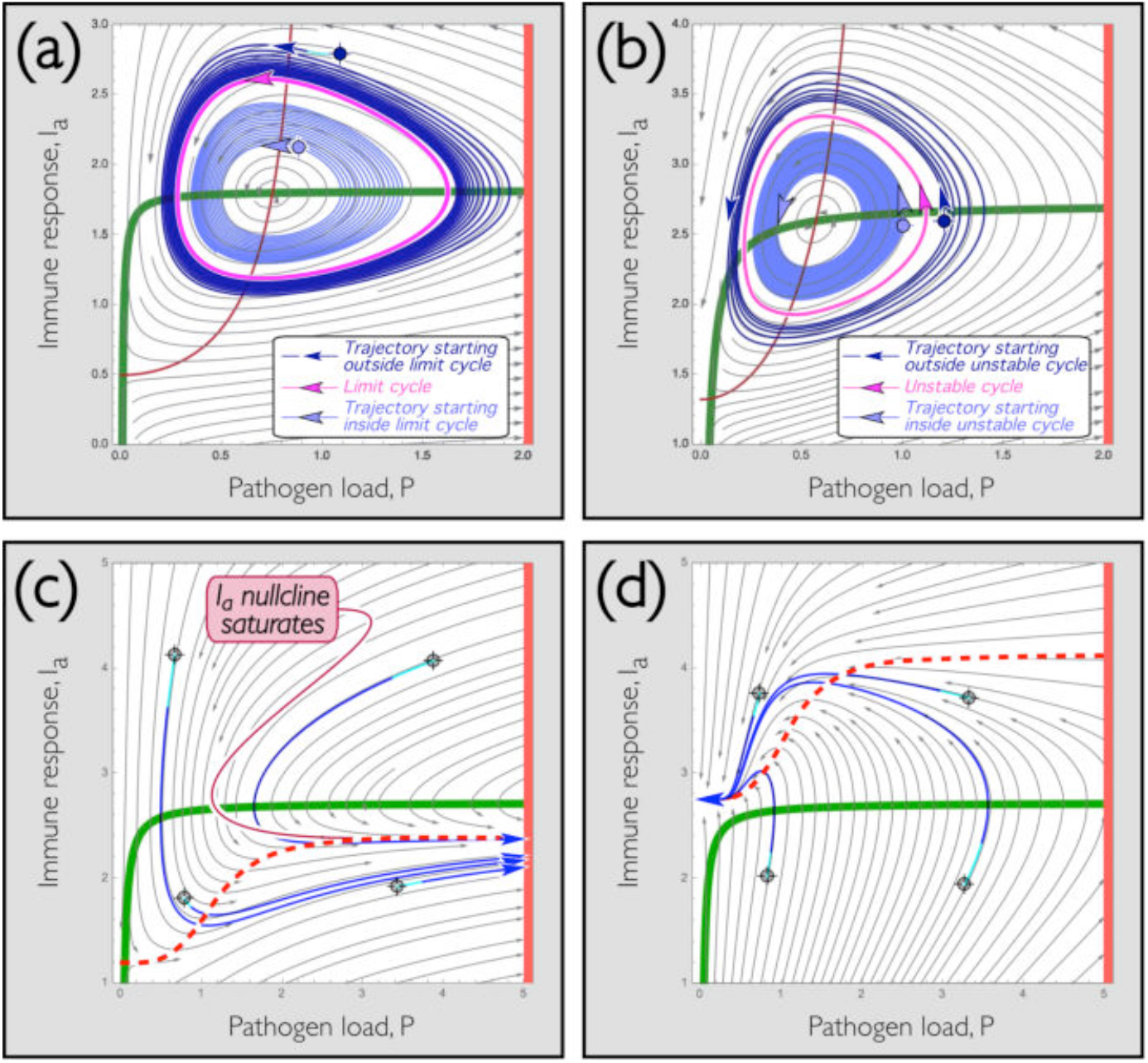
Cycles and bifurcations: In the presence of delay, multiple states are possible. **(a)** <u>Limit cycle</u>: When the central fixed point is unstable, trajectories nearby expand outward (light blue). When delay causes outer trajectories to become bounded (dark blue), a limiting cycle emerges (magenta). **(b)** <u>Unstable cycle</u>: The counterpoint to a limit cycle is an unstable cycle, shown here (magenta). Trajectories outside now expand outward (dark blue) and those inside contract inward (light blue). **(c)** <u>Tangent bifurcation to death</u>: By changing parameters, it is straightforward to produce a sigmoidal I_a_ nullcline that saturates at high pathogen load: red curve, dashed for emphasis. Reducing the setpoint and increasing the autoregulatory rate decreases the immune response (see text for values), causing the red and green nullclines to separate in a tangent bifurcation. Notice that all trajectories lead to death in this case. Notice ordinate is shifted upward compared with prior plots **(d)** <u>Tangent bifurcation to cure</u>: . Adjusting parameters to reduce the pathogen growth but strengthen the immune response (see text) causes the two nullclines to separate in a second tangent bifurcation. In this case, all initial states are cured.

Using a different initial state, we get an expanding trajectory, shown as light blue in Fig. 11(a). Since trajectories outside of the magenta orbit migrate inward, and trajectories inside migrate outward, it is a well-known consequence of the Poincaré-Bendixson theorem that the magenta orbit itself must be a limit cycle. That is, for long times, all trajectories within a bounded region outside of the dark blue trajectory must approach the magenta orbit – and correspondingly, **all states must periodically cycle indefinitely**. Technically, the guarantee of a limit cycle assumes that the system is two-dimensional; computationally this is close enough to being the case that a limit cycle does in fact appear.

For the same parameters as in Fig. 11(a), but using smaller delays than about *τ* = 0.1, the central fixed point becomes stable, and a constant (rather than periodic) chronic state is approached. For larger delays the fixed point becomes globally unstable, and either a cure or death is reached as in previous examples. We note in both Fig. 11(a) and 11(b), shown next, actual trajectories deviate from the streamlines shown in the background. This is a signature of the delay, which causes trajectories to follow derivatives obtained earlier in time than those produced by instantaneous evaluation.

Other states are also easily produced; for example in Fig. 11(b) we show the opposite of a limit cycle: an unstable cycle obtained at *g* = 3, *k* = 1.1, *k* = 0.08, *g* = 1, *f* = 0.5, *l* = 1.325 and *τ* = 0.1. Trajectories beginning outside of that cycle expand to either reach a cure or death, and trajectories inside of the cycle orbit repeatedly and slowly approach a stable chronic state. It is unclear if this type of behavior is seen physiologically: it would represent a **chronic state that, if significantly perturbed by treatment or environment, could either degenerate into a cure or death**: a worrying possibility!

Finally, we remark that other behaviors than those examined here are also possible. A couple of relatively simple examples that do not rely on delay can be generated by increasing the cooperativity, *n*, from 2 to 4. This makes the red nullcline strongly sigmoidal as shown in Fig’s 11(c) & (d). The nullcline is weakly sigmoidal with *n* = *2*, but *n* = 4 is preferable for illustrative purposes. We can lower the sigmoid on the *I*_*a*_ axis by changing *f*_*a*_ to 2 and *l*_w_ to 1.2 but keeping the other parameters as before (*g*_*p*_ = 3, *k*_*a*_ = 1.1, *k*_*i*_ = 0.08, *g*_*a*_ = 1 and *τ* = 0.1). This causes the two nullclines to separate as shown in Fig. 11(c). This separation, referred to as a tangent bifurcation, eliminates intersections between the nullclines, and so the fixed points at the intersections vanish, and so does the possibility of chronic or recurrent states. Rather, as the blue trajectories starting from four different initial states illustrate, **all trajectories end in death**. Biologically, this occurs because the lower immune setpoint (*l*_w_ → 1.*2* instead of 1.325) combined with the faster regulation to that setpoint (*f*_*a*_ → *2* instead of 0.5) causes the adaptive immune response to rapidly diminish as pathogen load increases. As shown in Fig. 11(c), this forces all trajectories to quickly approach the saturation value of the *I*_*a*_ nullcline (*I*_*a*_ = 1.38), irrespective of initial state.

The tangent bifurcation shown in Fig. 11(c) arose when the sigmoidal red curve was moved downward (by changing *f*_*a*_ and *l*_o_ at increased *n*). We can instead move that nullcline upward to produce a second tangent bifurcation, as shown in Fig. 11(d). To do this, we keep *n* = 4 and set *g*_*p*_ = 1, *f*_*a*_ = 3, *l*_o_ = *2*.75. The other parameters remain as before (*k*_*a*_ = 1.1, *k*_*i*_ = 0.08 and *τ* = 0.1). As in Fig. 11(c), these parameters are chosen for illustrative clarity, and are not special: many other values will also produce a sigmoidal *I*_*a*_ nullcline separated above from the green *P* nullcline. Whenever this separation occurs, **all initial states end in a cure**, as shown by the blue trajectories in the figure. This isn’t mysterious: biologically, it arises because the pathogen growth rate, *g*_*p*_, was decreased from 3 to 1, and immune values are consequently forced rapidly (*f*_*a*_ → 3 instead of *2*) to a high setpoint (*l*_w_ → *2*.75 instead of 1.*2*). This represents a slowly growing pathogen attacked by immune agents that rapidly grow to a high value. Naturally enough, such infections are invariably cured.

As we mentioned, the bifurcations shown in Fig. 11(c) & (d) do not rely on delay; nevertheless they hint at other possible behaviors. The Mackey-Glass^20^ equation defines a single variable that describes growth of blood cells, and when delay is included this produces chaotic and other complicated solutions. We have analyzed straightforward dynamics of two variables in our equations [6] and [10], and these dynamics already exhibit plentiful dynamical complexities, many of which duplicate behaviors known to be present in immune responses, and a few of which may represent new unreported states. It seems inevitable that more detailed analysis including delay is bound, like the simpler Mackey-Glass system on which our model is based, to exhibit even more complicated solutions.

### Conclusion

To Physicists, it is surprising that infections that grow exponentially require an exposure threshold to persist, and that once an infection that could be successfully treated by antibiotics exceeds a larger threshold, no amount of antibiotic will prevent its growth.

To Biologists, it is well known, but difficult to understand, why some infections persist, either in a relatively stable chronic form, or in a periodic state that regularly grows and shrinks. Likewise it is unexplained why some patients get worse before getting better, while others get better before getting worse. Nor why one patient can encounter a disease and survive, while a medically indistinguishable patient may succumb.

All of these observations are explained by a minimal mathematical model of competition between a pathogen and an immune system consisting of both innate and adaptive parts. Specifically, we have found the following.

First, considering the case where delay in mobilizing an adaptive immune response is neglected:

1. The innate immune system alone produces a **critical pathogen load**, *P*_*c*_ below which the pathogen will die out, and above which the pathogen will grow. This provides an explanation for why we can tolerate limited exposure to infections.
2. The adaptive system alone produces an attractive fixed point with constant pathogen load and immune response. This can be interpreted as a **chronic state**. Moreover, autoregulation of the adaptive system can lead to recurrent cycles of infection.
3. **Slow immunosuppression can reinforce these recurrent cycles**, while **fast immunosuppression** enlarges the growth and shrinkage of pathogen levels during these cycles, leading potentially to **dramatic pathogen growth**.
4. **Combining innate and adaptive immune responses allows more serious infections (that neither system alone could control) to be resolved**. In short, the innate system can clear a minor infection, but cannot cope with a larger infection. The adaptive system can stabilize a larger infection, but cannot by itself destroy the infection. Combined, the two systems can both stabilize and eliminate larger infections.
5. Infections can exhibit **several distinct types of dynamics**. They can monotonically improve or worsen. They can worsen and then improve. They can improve and then worsen. And they can cycle repeatedly before ultimately resulting in a cure or in death.
6. The states, defined by initial pathogen load and adaptive immune status, leading to ultimate cure or death can be very closely interleaved, especially close to the chronic steady state. This has two implications. First, **small differences in initial state can produce large differences in outcome**. Second, in principle intentional intervention could steer the competition between pathogen and immune system to a state leading to a successful outcome.
7. With respect to interventions:
  a. **Early intervention** (e.g. with antibiotics) can produce a cure that later intervention may not.
  b. Indeed, **once a pathogen level has exceeded a point of no return, no amount of intervention may effect a cure**.
  c. As one would expect, vaccination or prior infection greatly improves the response to a new pathogen exposure.
  Second, if delay of the adaptive system is included:
8. Paradoxical outcomes arise, for example **a patient subjected to a lower pathogen load may succumb while a comparable patient subjected to a larger load may survive**. This appears to be due to the ability of a weaker pathogen load to grow to a dangerous level in the presence of a weaker immune system. This permits the pathogen to sneak up on the immune system, before the immune system has mobilized an effective response. The nonlinearity of the adaptive system produces a significantly stronger response to higher pathogen loads, which can produce the paradoxical outcome in which a sicker patient may survive while a less sick one expires.
9. **These paradoxical responses appear to be generic** in that simpler as well as more realistic delay models can both generate alternating cure and death outcomes as the pathogen load is increased.
10. Other, more complicated states are also possible, including stable limit cycles and chronic states that can degenerate into a cure or death.

These are purely mathematical findings that follow from a straightforward model that utilizes a minimum of assumptions about the detailed workings of the immune system. We stress that our mathematical results can absolutely not be taken to suggest that more detailed and realistic workings of the immune system and of pathogenic evasion strategies are not important. Rather, our findings show that mathematics, on its own and without those details, provides mechanisms for a number of veridical dynamical behaviors that may otherwise be difficult to rationalize. Other possible complicated dynamics are also present in this system, including stable limit cycles and chronic states that can degenerate into a cure or death. Based on prior work of related equations, more complicated behaviors also seem likely.

This work is partially supported by NSF CBET, award no. 1804286.

The data that supports the findings of this study are available within the article.

## Notes

### Competing Interest Statement

The authors have declared no competing interest.

